# Identification of plasmid-mediated tigecycline resistance *tet*(x4) and New Delhi Metallo-β-lactamase (NDM) in an *Escherichia coli* isolate from Canada

**DOI:** 10.1101/2025.03.27.645688

**Authors:** Laura F. Mataseje, Nicole Lerminiaux, Allison McGeer, Jennie Johnstone, Susan Poutanen, Erin McGill, Yves Longtin, the Canadian Nosocomial Infection Surveillance Program (CNISP)

## Abstract

Rapid spread of carbapenemase producing Enterobacterales is a global health threat for which Tigecycline is one of the few remaining therapeutic options. Resistance by *tet*(x4) is particularly worrisome given its presence on mobile genetic elements. Here we report on a tigecycline-resistant *E*.*coli* strain isolated from a patient returning from Pakistan. A *tet*(x4) gene was found on an IncFII plasmid isolated from an ST410 *E*.*coli* co-harbouring NDM-5. The combination of carbapenemase and *tet*(x4) is concerning.

## Brief Report

### Introduction, Materials and Methods, Results/Discussion

The rapid increase of extensively drug-resistant (XDR) Gram-negative bacteria, particularly carbapenem-resistant Enterobacterales, has made treatment challenging. Tigecycline is one of the few last resort therapeutic options available for treating multidrug-resistant Gram-negative bacterial infections ^1^. However, the newly emerged plasmid-encoded *tet*(X) and efflux *tmex*CD1-*topr*J1 genes in Enterobacterales encode high level tigecycline resistance (TIG-R) and pose a threat to its clinical efficacy ^2,3^. Plasmid-mediated resistance conferred by *tet*(x) genes was first reported in 2019 from *Acinetobacter* and Enterobacterales of both human and animal origin ^3^.

This publication included the first report of *tet*(x4), which was shown to inactivate all tetracyclines, including tigecycline, eravacycline and omadacycline. It was isolated from *E. coli* of clinical origin co-harbouring NDM ^3^. Since then, a few additional reports have identified *tet*(x4) co-harboured with a carbapenemase and/or mobile colistin resistant genes (mcr-type) ^4–8^. Here we report the first case of a plasmid mediated *tet*(x4) co-located in an NDM-producing *E*.*coli* isolated from a patient in Canada.

In 2021, a 79 year old male with diabetes was hospitalized in Canada. The patient had recently traveled to Pakistan for a medical procedure. Routine admission screening for carbapenemase-producing organisms was performed by direct plating of a pooled nasal/rectal swab onto ESBL isolation media (Oxoid), followed by meropenem disc diffusion screening and a β-CARB test (Biorad). An *E. coli* isolate confirmed by PCR (Xpert CarbaR, Cepheid) to harbor NDM and OXA-48-type was recovered and sent to the National Microbiology Laboratory, where antimicrobial susceptibility testing by Sensititre™ and whole genome sequencing using both the Illumina NextSeq Platform (Illumina, San Diego, USA) and Oxford Nanopore Technologies (ONT) (Oxford, Oxfordshire, UK) Minion platform were performed. Short-read libraries were created with TruSeq Nano DNA HT sample preparation kits (Illumina) and run on an Illumina NextSeq™ platform. Illumina reads had adaptors trimmed and were filtered for an average Q-score > 30 with trim-galore v0.6.7 (https://github.com/FelixKrueger/TrimGalore). Long-read sequences were generated using the Rapid Barcoding Kit (SQK-RBK004) on R9.4.1 flow cells. Read data was basecalled and demultiplexed with Guppy v6.5.7 using the Super High Accuracy model (ONT). Hybrid assemblies were generated using Unicycler v0.5.0 (https://github.com/rrwick/Unicycler) using default settings. Illumina sequence data and hybrid assemblies are deposited under Bioproject PRJNA1240299. StarAMR v0.10.0 ^9^ was used to detect antimicrobial resistance genes, plasmid replicons and sequence types, and MOB-suite v3.1.4 (https://github.com/phac-nml/mob-suite) was used to assess plasmid mobility. The original culture was mixed with two colony morphologies. Two carbapenemase producing *E*.*coli* isolates were recovered: A21005 harboring OXA-48-type and NDM and A21005-2 harboring NDM. Susceptibility data showed near identical profiles with the exception of TIG-R in A21005-2 (Table 1). Both isolates were non-susceptible to all drugs tested with the exception of aminoglycosides, plazomicin, fosfomycin and cefiderocol, making both isolates XDR ^10^.

**Table 1.**
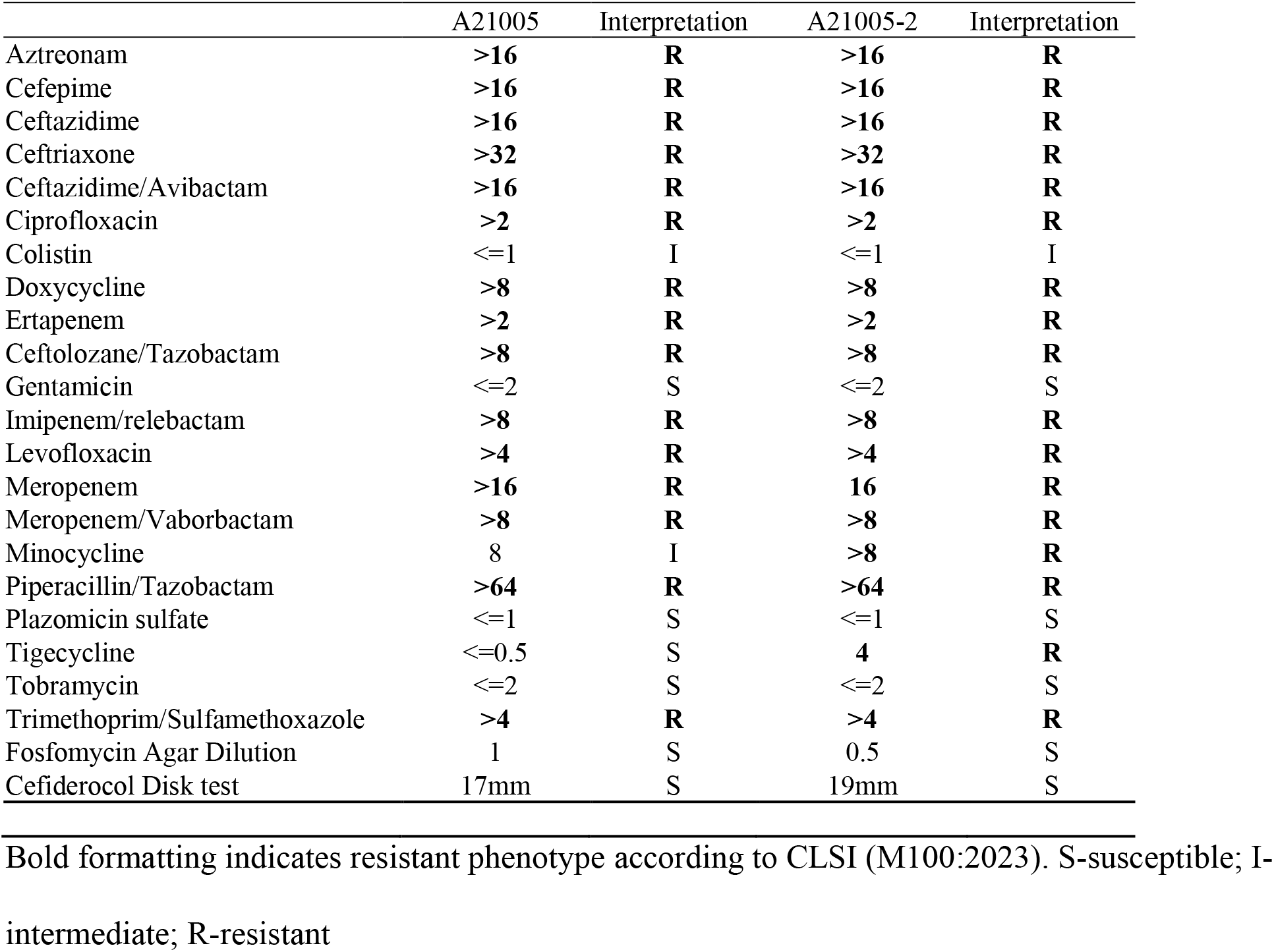
Antimicrobial susceptibilities (mg/L) for strains tested in this report determined using Sensititre panel CAN1MSTF.

|  | A21005 | Interpretation | A21005-2 | Interpretation |
| --- | --- | --- | --- | --- |
| Aztreonam | >**16** | **R** | >**16** | **R** |
| Cefepime | >**16** | **R** | >**16** | **R** |
| Ceftazidime | >**16** | **R** | >**16** | **R** |
| Ceftriaxone | >**32** | **R** | >**32** | **R** |
| Ceftazidime/Avibactam | >**16** | **R** | >**16** | **R** |
| Ciprofloxacin | >**2** | **R** | >**2** | **R** |
| Colistin | <=1 | I | <=1 | I |
| Doxycycline | >**8** | **R** | >**8** | **R** |
| Ertapenem | >**2** | **R** | >**2** | **R** |
| Ceftolozane/Tazobactam | >**8** | **R** | >**8** | **R** |
| Gentamicin | <=2 | S | <=2 | S |
| Imipenem/relebactam | >**8** | **R** | >**8** | **R** |
| Levofloxacin | >**4** | **R** | >**4** | **R** |
| Meropenem | >**16** | **R** | **16** | **R** |
| Meropenem/Vaborbactam | >**8** | **R** | >**8** | **R** |
| Minocycline | 8 | I | >**8** | **R** |
| Piperacillin/Tazobactam | >**64** | **R** | >**64** | **R** |
| Plazomicin sulfate | <=1 | S | <=1 | S |
| Tigecycline | <=0.5 | S | **4** | **R** |
| Tobramycin | <=2 | S | <=2 | S |
| Trimethoprim/Sulfamethoxazole | >**4** | **R** | >**4** | **R** |
| Fosfomycin Agar Dilution | 1 | S | 0.5 | S |
| Cefiderocol Disk test | 17mm | S | 19mm | S |

Bold formatting indicates resistant phenotype according to CLSI (M100:2023). S-susceptible; I- intermediate; R-resistant

Complete closed genomes and plasmids were used to examine AMR genes and plasmids between the two *E*.*coli* (Table 2). The first isolate (A21005) belonged to the high risk clone sequence type (ST) 361. It contained seven plasmids ranging in size from 6-176Kb. Of note, it contained two different *bla*_CTX-M-15_ plasmids, a *bla*_CMY-42_ plasmid, a *bla*_OXA-232_ plasmid and a *bla*_NDM-5_ plasmid. The second case (A21005-2) belonged to ST410 and harboured four different plasmids, of note, a *bla*_CMY-146_, *bla*_NDM-5_ and *tet*(x4) plasmid were observed. Both isolates contained similar plasmid content including a non-AMR 176kB plasmid (repA(pKOX)), however, the sizes of all other plasmids between the two isolates of the same replicon differed. In both isolates, the *bla*_NDM-5_ gene was found on an IncF-type plasmid, but these plasmids do not appear to be related (<50% identity) and are likely separate acquisitions (data not shown). *E*.*coli* ST410 is known as a high risk clone and has been reported worldwide ^11^. The ST410 lineage has been shown to cause recurrent infections in humans, as well as hospital outbreaks ^11^.

**Table 2.** Summary of genotypic data on tetracycline resistant (A21005-2) and tetracycline sensitive (A21005) cases. Total content is shown for each isolate followed by each plasmid.

| Isolate | Plasmid name | AMR genes | Plasmid | Sequence Type | Plasmid Size |
| --- | --- | --- | --- | --- | --- |
| A21005 |  | aadA2, aadA5, blaCMY-42, blaCTX-M-15, blaCTX-M-15, blaNDM-5, blaOXA-232, blaTEM-1B, dfrA12, dfrA17, mph(A), qacE, qacE, qnrS1, sul1, sul1, tet(B) | ColKP3, IncFIB(pB171), IncFII, IncI(Gamma), IncY, repA(pKOX) | 361 | n/a |
|  | pA21005_A | none | repA(pKOX) | n/a | 176160 |
|  | pA21005_B | blaCTX-M-15, qnrS1 | unknown | n/a | 91863 |
|  | pA21005_C | none | IncY | n/a | 89633 |
|  | pA21005_D | aadA5, blaCTX-M-15, blaTEM-1B, dfrA17, mph(A), qacE, sul1, tet(B) | IncFIB(pB171) | n/a | 87008 |
|  | pA21005_E | aadA2, blaNDM-5, dfrA12, qacE, sul1 | IncFII | n/a | 82339 |
|  | pA21005_F | blaCMY-42 | IncI(Gamma) | n/a | 47019 |
|  | pA21005_G | blaOXA-232 | ColKP3 | n/a | 6141 |
| A21005-2 |  | aadA2, blaCMY-146, blaNDM-5, blaTEM-215, blaTEM-215, dfrA12, floR, mph(A), qacE, qepA4, sul1, tet(X4) | IncFIB(pB171), IncFII, IncI(Gamma), repA(pKOX) | 410 | n/a |
|  | pA21005-2_A | none | repA(pKOX) | n/a | 176160 |
|  | pA21005-2_B | aadA2, blaNDM-5, blaTEM-215, dfrA12, mph(A), qacE, qepA4, sul1 | IncFIB(pB171) | n/a | 93954 |
|  | pA21005-2_C | blaTEM-215, floR, tet(X4) | IncFII | n/a | 87663 |
|  | pA21005-2_D | blaCMY-146 | IncI(Gamma) | n/a | 24054 |

The *tet*(x4) gene was harboured on a 87Kb IncFII plasmid (p0821005-2_C) that also contained the resistance genes *flo*R and *bla*_TEM-125_. (Figure 1). This plasmid was predicted to be conjugative by MOB-suite with a MOBF relaxase and MPF_F mating pair formation protein. This plasmid encoded the *hok-sok* Type I toxin-antitoxin system, antirestriction protein KlcA, single-stranded binding protein Ssb, an SOS inhibitor protein PsiB, and several genes involved in plasmid stability (*stbB* and *parM*). Similar plasmids have been observed from *E*.*coli* isolated from chickens in Pakistan ^8^ as shown in the Figure. Other studies have reported the *tet*(x4) gene on various other plasmid backbones including IncX ^4,12^, IncQ ^5,6^ and hybrid plasmids ^4,12^. Though some studies have characterized the genetic environment by Types (I and II) ^12^ or by Groups (G1-G4) ^4^ the complete mobile elements that carries *tet*(x4) has not been fully described. Here we observed Tn3-IS*26*-IS*CR*-*tet*(x4)-est*T*-IS*CR*-*floR*-*virD2*-IS*26*-*bla*TEM-215-IS*26*-*ramA-*Tn3 among several hypothetic open reading frames as well as a DNA-methyltransferase and a restriction endonuclease (DEAD/DEAH). The duplicated Tn3 was flanked by 38bp inverted repeats and a 5bp target site duplication making the total size 28 198bp (Figure 1). Though the *E*.*coli* ST361 also harboured an IncFII plasmid these were only 70% similar to each other and didn’t share similar AMR genes.

**Figure 1.**
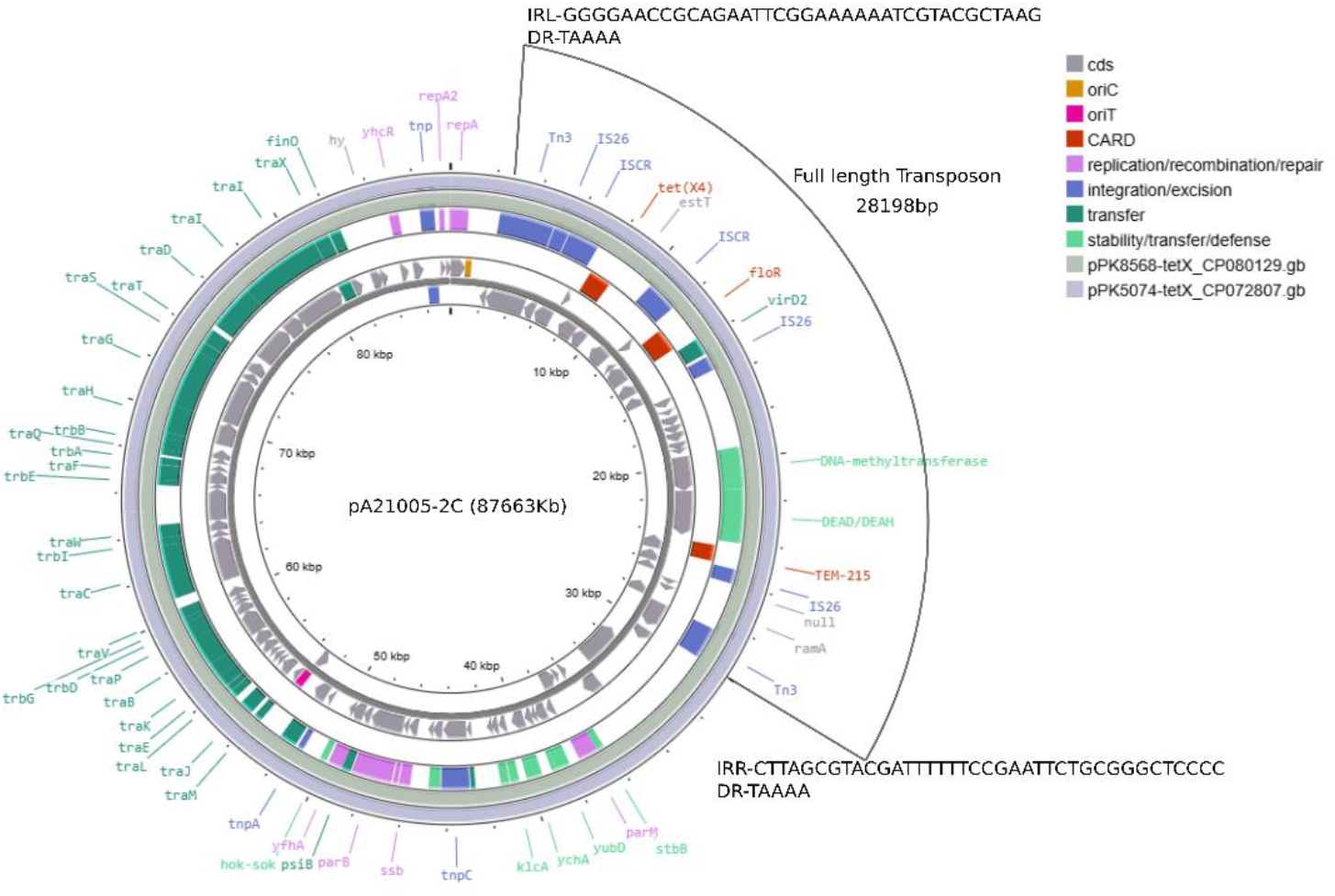
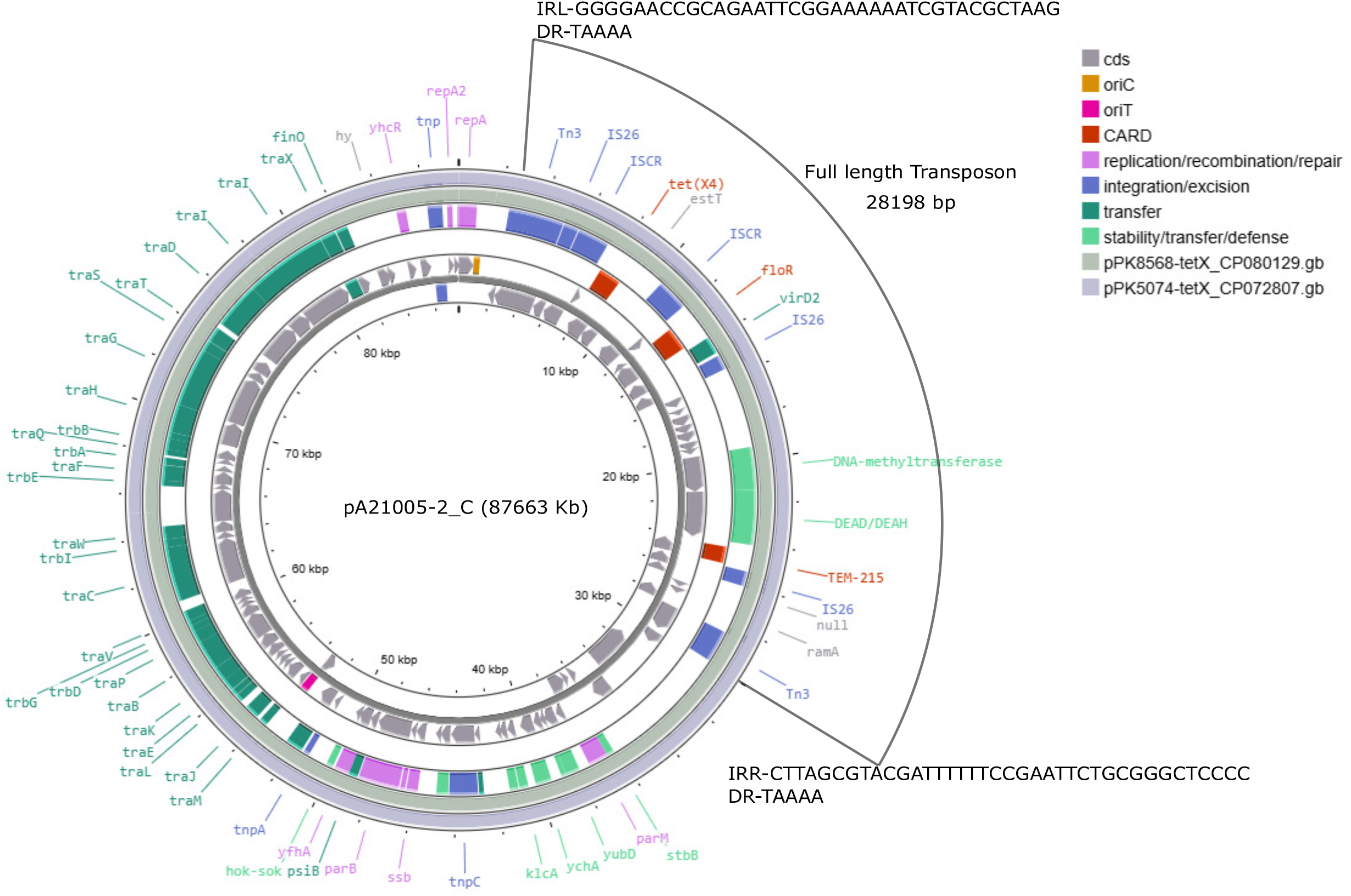
Plasmid generated with Proksee. Innermost two tracks represent forward and reverse open reading frames of pA21005-2. Middle two tracks are annotations generated by CARD RGI and mobileOG databases. Outer two tracks are comparisons to other *tet*(x) plasmids using BLAST.

In summary, we observed two *E. coli* isolates from the same patient that were unrelated clonally or by AMR plasmid transfer. Both were high risk global clones and harboured multiple AMR plasmids including carbapenemases overall making them XDR. To the authors’ knowledge this is the first report of a plasmid-mediated tigecycline resistance gene detected in Canada. Its occurrence on a plasmid with other resistance genes, co-harboured with a carbapenemase and with a high-risk global clone is of significant concern.

## Acknowledgments

We would like to thank Ken Fakharuddin for technical work and The Canadian Nosocomial Infection Surveillance Program (CNISP) carbapenemase producing organisms working group members: We thank the physicians, epidemiologists, infection control practitioners, and laboratory staff at each participating hospital for their contributions to the study and would like to recognize the CNISP CPO Working Group: Annie-Kim Nguyen (Public Health Agency of Canada, Montreal, Quebec), Robyn Mitchel (Public Health Agency of Canada, Ottawa, Ontario), Kevin Katz (North York General Hospital, Toronto, Ontario), Kris Cannon (Alberta Health Services, Calgary, Alberta), Ian Davis (QEII Health Sciences Centre, Halifax, Nova Scotia), Tamara Duncombe (Fraser Health Authority, Vancouver, British Columbia), Chelsey Ellis (The Moncton Hospital, Moncton, New Brunswick), Jennifer Ellison (Alberta Health Services, Calgary, Alberta), Aleks Gara (Vancouver Coastal Health, Vancouver, British Columbia), Jennifer Happe (Alberta Children’s Hospital, Calgary, Alberta), Susy Hota (University Health Network, Toronto, Ontario), Pamela Kibsey (Royal Jubilee Hospital, Victoria, British Columbia), Santina Lee (University of Manitoba Children’s Hospital, Winnipeg, Manitoba), Jerome A. Leis (Sunnybrook Health Sciences Centre, Toronto, Ontario), Xena Li (North York General Hospital, Toronto, Ontario), Shazia Masud (Fraser Health Authority, Vancouver, British Columbia), Allison McGeer (Sinai Health, Toronto, Ontario), Jessica Minion (Saskatchewan Health Authority, Regina, Saskatchewan), Sonja Musto (Health Sciences Centre, Winnipeg, Manitoba), Kishori Naik (West Park Health Care Centre, Toronto, Ontario), Senthuri Paramalingam (Birchmount and Centenary Hospitals, Toronto, Ontario), Connie Patterson (McGill University Health Centre, Montréal, Québec), Nancy Petitclerc (Hôpital Maisonneuve-Rosemont, Montréal, Québec), Ewa Rajda (McGill University Health Centre, Montréal, Québec), Katy Short (Fraser Health Authority, Vancouver, British Columbia), Stephanie W. Smith (University of Alberta Hospital, Edmonton, Alberta), Jocelyn A. Srigley (BC Women’s and BC Children’s Hospital, Vancouver, British Columbia), Kathy N Suh (The Ottawa Hospital, Ottawa, Ontario), Dharma Teja Yalamanchili (Saskatchewan Health Authority, Regina, Saskatchewan) Nisha Thampi (Children’s Hospital of Eastern Ontario, Ottawa, Ontario), Reena Titoria (Provincial Health Services Authority, Vancouver, British Columbia), Jen Tomlinson (Health Sciences Centre, Winnipeg, Manitoba), Joseph Vayalumkal (Alberta Children’s Hospital, Calgary, Alberta), Kristen Versluys (Alberta Health Services, Edmonton, Alberta), Diana Whellams (Fraser Health Authority, Vancouver, British Columbia), Titus Wong (Vancouver Coastal Health Research Institute, Vancouver, British Columbia).

